# Immune dynamics at single cell protein level after delta/omicron infection in COVID-19 vaccinated convalescent individuals

**DOI:** 10.1101/2022.09.05.506626

**Authors:** Rimpi Bajaj, Zhiqi Yang, Vincent Hammer, Simone Pöschel, Kristin Bieber, Madhuri S Salker, Nicolas Casadei, Stephan Ossowski, Olaf Riess, Yogesh Singh

## Abstract

Both COVID-19 mRNA or recombinant Adenovirus vector (rAdVV) based vaccines have shown a great efficacy in generating humoral and cellular immune responses. Two doses of the COVID-19 vaccines generate enough antibodies and generate spike-specific T cell responses. However, after 6-8 months there is a decline in antibody production and T cell responses. Due to the rise of new SARS-CoV-2 variants of concern, a third or even fourth dose of vaccine was recommended for the elderly, immune comprised and frontline medical health care workers. However, despite additional booster doses given, those who were infected with either delta or omicron (during December 2021 – March 2022) had symptoms of illness. By what means these COVID-19 vaccines provide immunity against the SARS-CoV-2 virus at the molecular level is not explored extensively yet and, it is an emerging research field as to how the SARS-CoV-2 virus is able to evade the host immunity. Most of the infected people had mild symptoms whilst some were asymptomatic. Many of the people had developed nucleocapsid antibodies against the SARS-CoV-2 delta/omicron variants confirming a humoral immune response against viral infection. Furthermore, cellular analysis shows that post-vaccinated recovered COVID-19 individuals have significantly reduced NK cells and increased T naïve CD4^+^, TEM CD8^+^ and B cells. This decrease in cellular immunity corresponds to individuals who recovered from alpha variants infection and had mild symptoms. Our results highlight that booster doses clearly reduce the severity of infection against delta/omicron infection. Furthermore, our cellular and humoral immune system is trained by vaccines and ready to deal with breakthrough infections in the future.

## Introduction

During the first stage of the pandemic, the virus remained genetically and antigenically stable. However, during the second stage of pandemic with significant genetic changes started to occur in the spike gene of the SARS-CoV-2 virus compared to the original Wuhan strain (Wild Type strain)^1^. Furthermore, as the virus continues to evolve, genetic mutations have been found in the functional domain of Spike (S) protein of SARS-CoV-2, which results in the altered phenotypes of the virus^2^. So far, the World Health Organization (WHO) has described five variants of concern (VOC) based on the changes in COVID-19 epidemiology, increase in pathogenicity, possibility of increase in transmissibility, disease presentation, vaccines, and treatment^3^. These VOCs elaborated B.1.1.7 (alpha), B.1.351 (beta), B.1.617.2 (delta), B.1.1.28 (gamma), and B.1.1.529 (omicron). Since November 2021, omicron has spread rapidly and has caused more infections than the original WT strain (Wuhan strain) and other strains^4^. Further research is required to understand the severity profile of omicron and the current evidence suggests that omicron severity is not high compared with other VOC^5^. However, new sublineages of omicron – BA.2.12.1, BA.4 and BA.5 are surging globally and based on structural comparisons of the spike protein BA.4 and BA.5 display comparable binding affinity to BA.2.12.1 to angiotensin-converting enzyme 2 (ACE2)^6^. This reduction in severity could be reduced by vaccination and/or immunity developed from previous infections. Fast decline of spike S1 specific antibody levels have been observed in vaccinated individuals after 6-8 months^7^. Nonetheless, omicron variants are definitively escaping the immune response generated against COVID-19 vaccines as well as natural COVID-19 immunity.

In the EU, six vaccines have been evaluated and are currently being deployed by the European Medicine Agency (EMA); adenovirus vector-based Vaxzevria (ChAdOx1-S [AstraZeneca], JNJ-78436735 or Jcovden (Ad26.COV-S recombinant [Janssen]), mRNA-based Spikevax (mRNA-1273 [Moderna], Comirnaty BNT162b2 [Pfizer/BioNTech]), COVID-19 Vaccine Valneva (inactivated adjuvanted [Valneva GmbH]) and the protein-based Nuvaxovid (NVX-CoV2373 [Novavax]^8-10^. All these vaccines have been designed according to the Spike protein sequence from of original strain of SARS-CoV-2. These vaccines induce a robust immune response and provide protection against severe disease ^10,11^. After taking the first dose of vaccination, either mRNA or vector-based vaccines, spike S1 and receptor binding domain (RBD) IgG antibodies are developed and are somehow protected against infection and had less adverse side effects^12^. Most of the reports observed that mRNA vaccines induce robust cytotoxic CD8^+^ T cell response (T cell immunity) and an effective humoral response which means the generation of high affinity neutralizing antibodies, that are indicators of protective immunity after taking the second dose^13^. This protection may last up to 6-8 months after 2 doses of the vaccine. However, IgG antibody titres start to wane after 8 months ^7,14^.

An earlier study suggested that cross neutralization of beta and delta variants was appreciated in all vaccinated individuals, but neutralization antibody against omicron was much lower^15^. Furthermore, SARS-CoV-2-specific T cells were identified up to 6 months after vaccination and no appreciable differences were noticed among WT and variant-specific CD4^+^ or CD8^+^ T cell responses, including omicron. These data indicate minimal escape at the T cell level and potentially highlights the lack of neutralizing antibodies in limiting severe COVID-19^15^. Two recent published studies point to the importance of T cell immunity – one indicated that T cell immunity is preserved in some but not all omicron infected induviduals^16^ whilst the other suggested that no change in T cell immunity was found after omicron infection compared to triple vaccinated individuals^17^. Therefore, we sought to explore why there was reduced neutralising antibody response and in parallel no change in T cell response against omicron (re)infection in vaccinated individuals. We hypothesized, despite existing immunity generated via vaccination, the omicron variant could lead to an immune escape by evading the immune system and changing the effector functions.

To understand, the immune escape and memory response, we took a candidate approach and performed a comprehensive analysis of humoral and cellular immune response. Samples were taken from individuals who became infected after taking two - four doses of (mRNA or rAdVV-based) vaccinations. Firstly, we confirmed the viral infection in vaccinated individuals after recovery of infection and measured their antibody (IgG and IgG neutralizing antibody specific to VOC) levels. Next, we examined comprehensively innate and adaptive immune response in the recovered vaccinated individuals. We identified that humoral and cellular immunity was intact and omicron infection leads to reduced lymphocytes and increased monocytes as WT strain of COVID-19 virus.

## Results

### Altered humoral and cellular immune profiles COVID-19 post-vaccination (2^nd^, 3^rd^ and 4^th^) individuals recovered after delta/omicron infection

During our longitudinal COVID-19 vaccine study, we collected samples from 110 vaccinated individuals and followed them for 1 year which included 3^rd^ or 4^th^ booster doses (Singh Y et al., 2022, bioRxiv). In this cohort study (May 2021 – June 2022), we found that out of 110 participants, 32 vaccinated (29%) individuals were infected SARS-CoV-2 and approximately 23.6% had mild COVID-19 symptoms including cough, fever, sore throat and runny nose and roughly 5.4% were asymptomatic (**Table 1**). Blood samples were collected 14 – 30 days of infection (confirmed with RT-PCR). We isolated peripheral blood mononuclear cells (PBMCs) from participants who were infected after taking either 2^nd^, 3^rd,^ or 4^th^ dose of mRNA/AdVV based COVID-19 vaccinations (PoV2/3/4) but were recovered ‘vaccinated infected but recovered’ (VibR). We first performed an antibody binding, neutralizing antibody, T cell response assay and comprehensive immunophenotyping of VibR group and compared with post-booster dose samples. (**Fig. 1a**).

**Table1:**
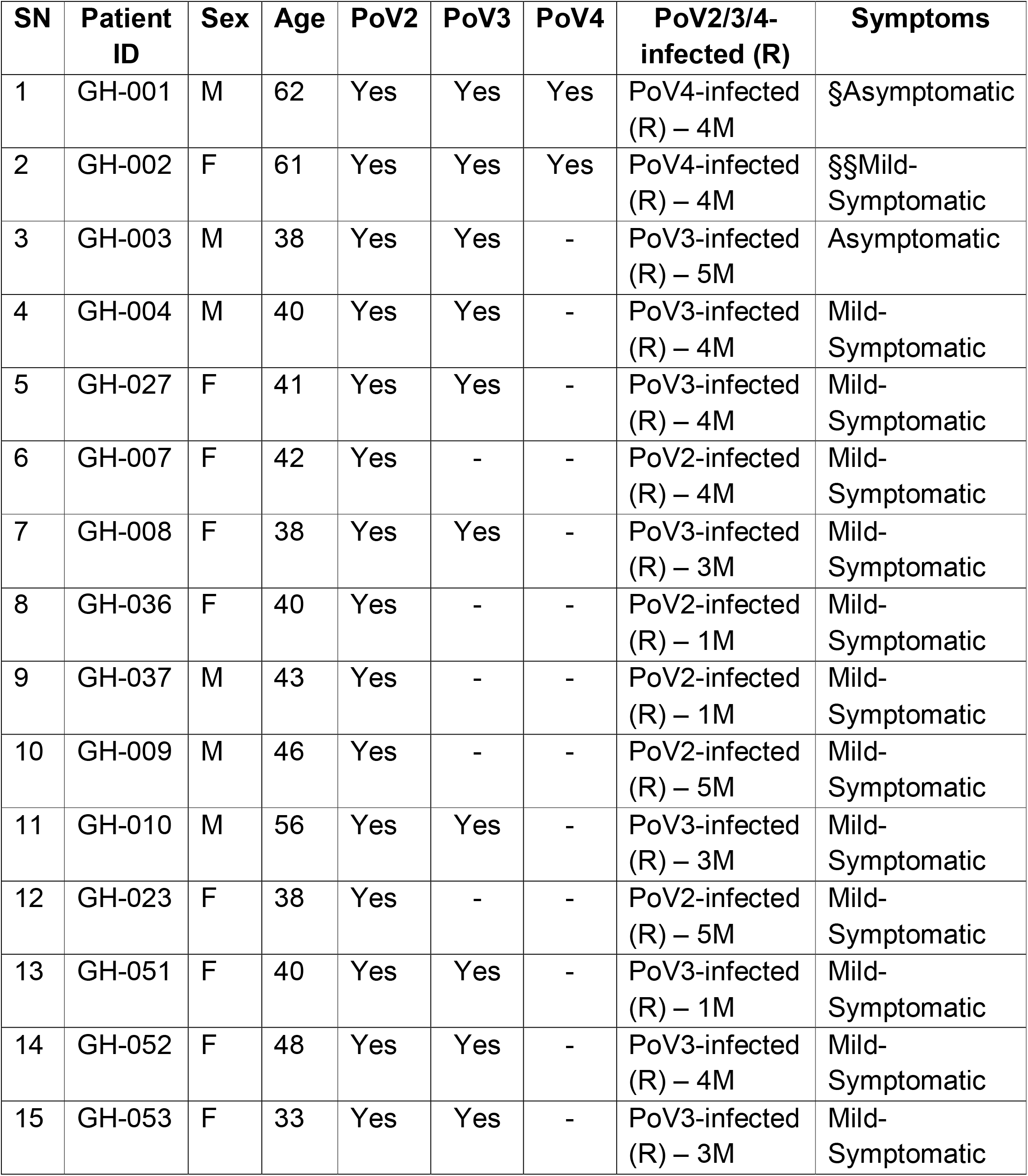
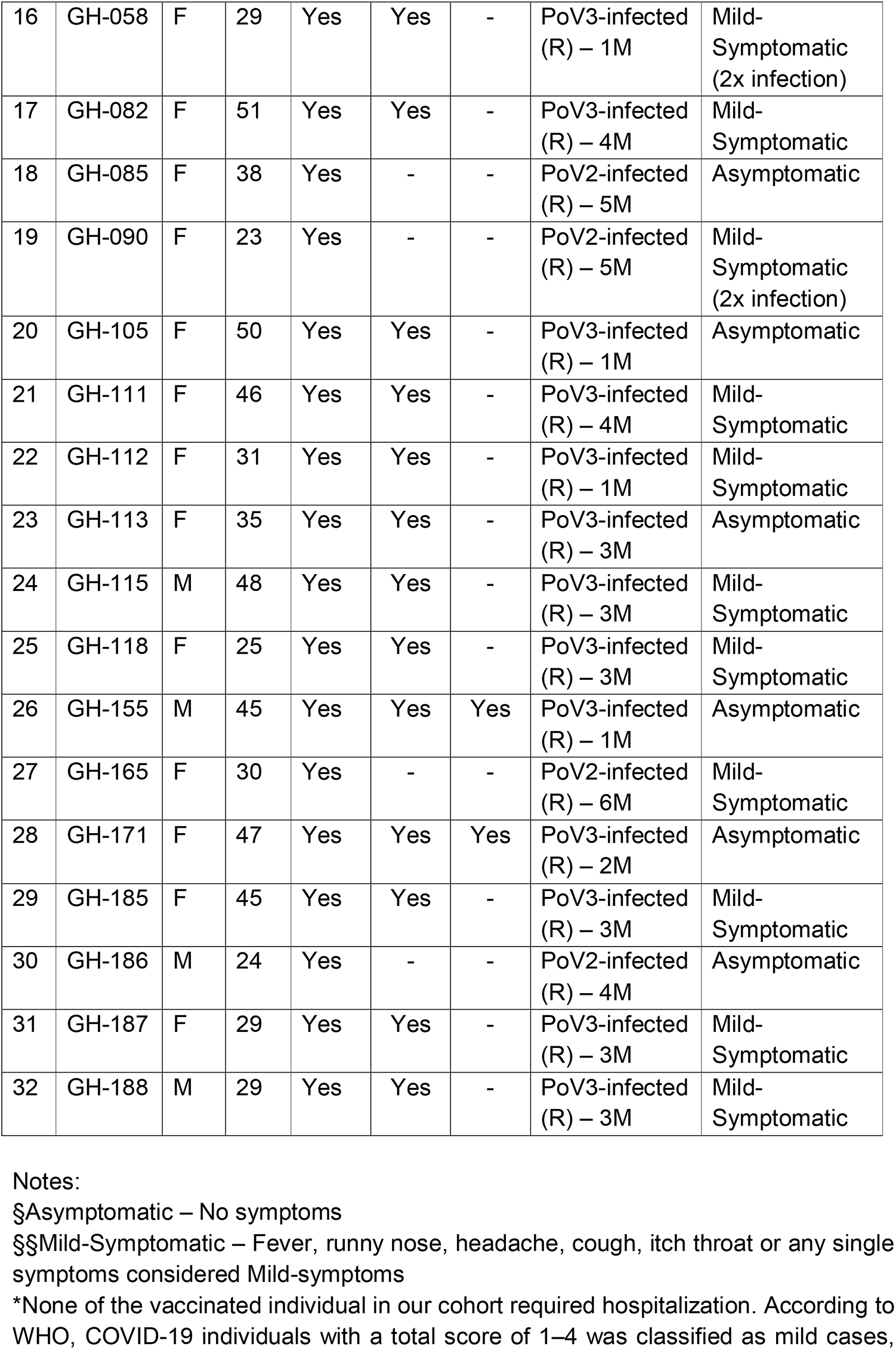
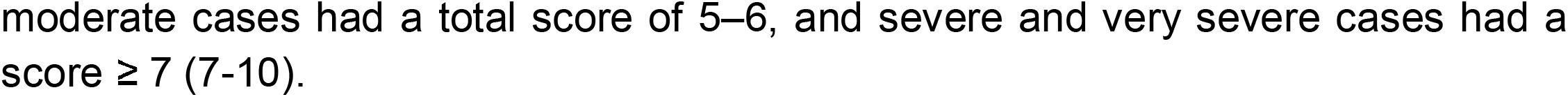
Patient demographics.

**Table 2:**
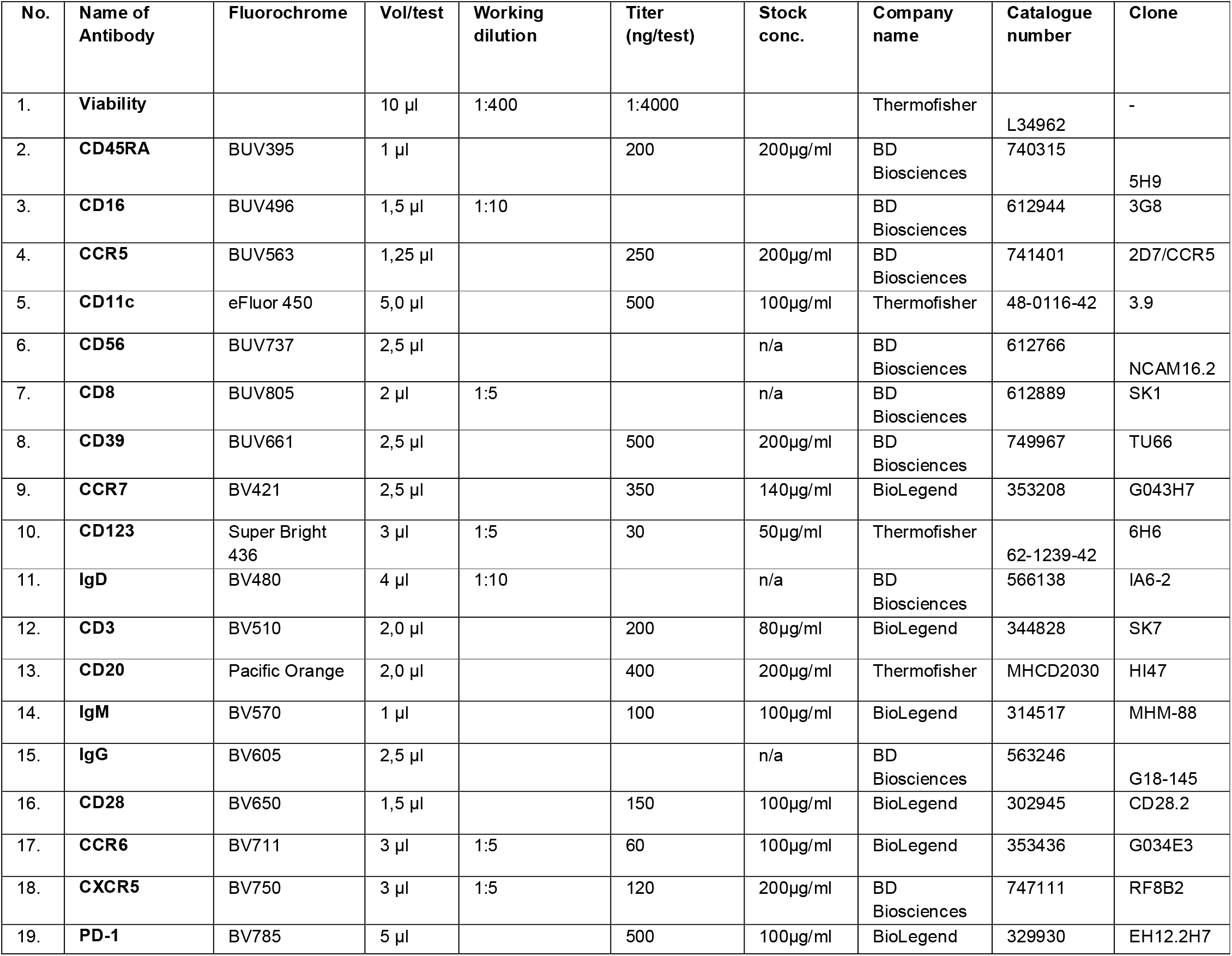
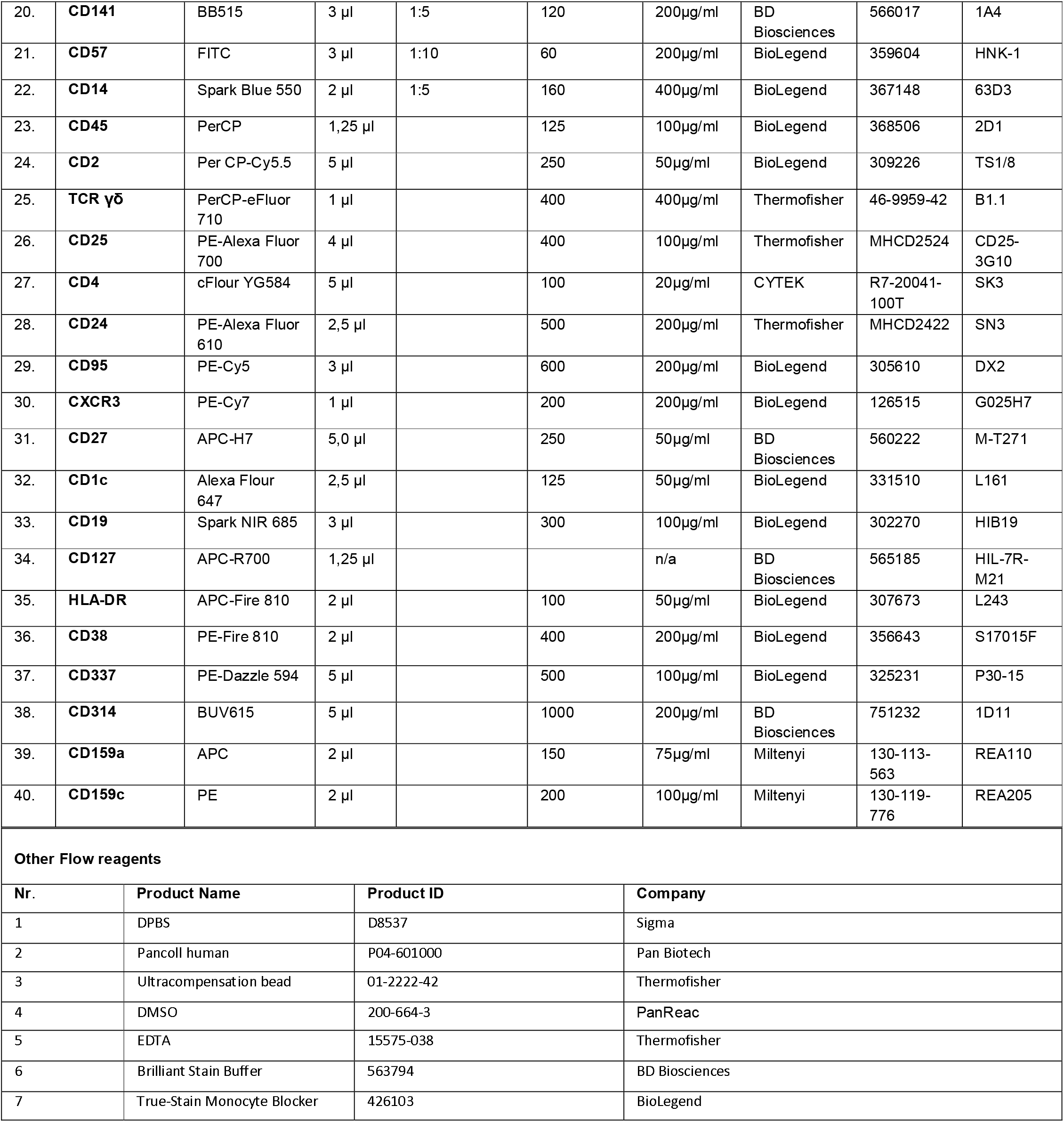
Antibodies used for staining 1-3×10^6^ PBMCs (fresh or frozen)/test.

**Fig. 1.**
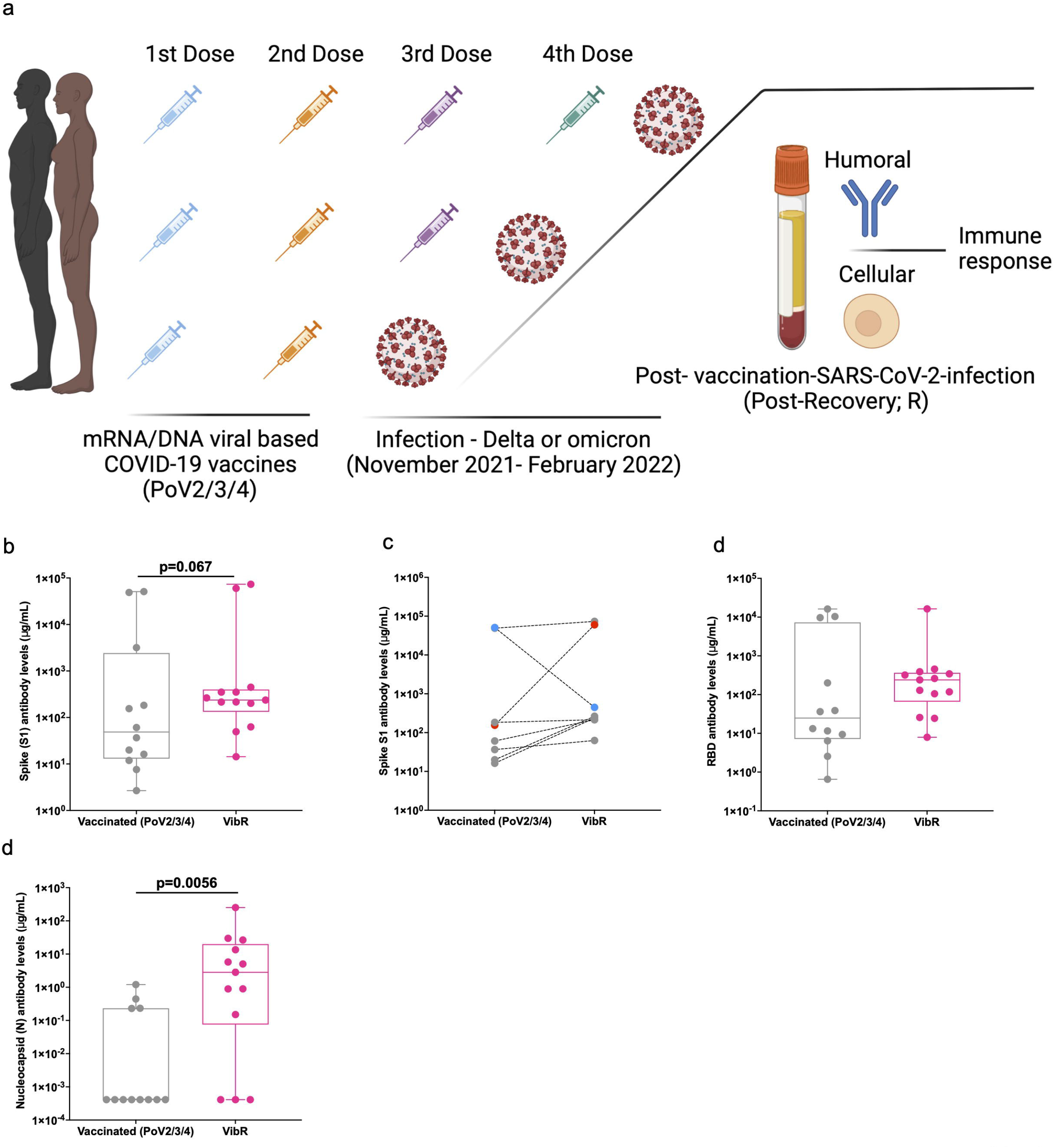
Study design and detection of virus specific binding antibodies before and after infection. (**a**) Schematic diagram representing the order of taking vaccination PoV2/3/4 followed by the approximate month scale of infection PoV-infection (VibR group). At post recovery, individuals provided the blood sample, for analysing plasma-specific to full length spike, RBD and Nucleocapsid proteins measured by ELISA and T cell response (**b**) Levels of binding S-specific antibodies before and after vaccination or infection. (**c**) RBD-specific antibody serum level measured by ELISA. (**d**) Nucleocapsid (N) antibody level detected before and after vaccination. Wilcoxon rank-sum test (unpaired) and Student T-test (paired samples) for two sample comparisons for PoV2/3/4 vs VibR groups. Violin plots indicate mean and frequency distribution of the data including the interquartile range. The exact P value is shown on the figures.

### Robust humoral immune response following breakthrough infection (post-recovery) after mRNA or DNA-based COVID-19 vaccines

There is a strong humoral immune response after SARS-CoV-2 infection in naïve people (>95%) and lasts for up to 6-8 months^18^, however some of them also do not develop humoral immune response^19^. Therefore, we first evaluated the humoral immune response by means of SARS-CoV-2 antibody response in plasma for spike S1 protein (S1), receptor binding domain (RBD) and nucleocapsid (N) using IgG-specific 3-plex antibody assay. Spike S1 protein specific assay against SARS-CoV-2 shows increase in antibody levels, however, it almost reached statistically significant different among in post recovered individuals compared to vaccinated individuals (**Fig. 1b**). A similar trend was seen in pair-wise comparison in eight of the individuals before (PoV2/3/4) and after infection (VibR group). One of the participants (red) showed an unusually higher level of S1-specific antibody after infection. In contrast, one (blue) showed dramatically lower antibody levels even after vaccination (**Fig. 1c**). Otherwise, most of the individuals show slightly increased level changes after infection (**Fig. 1c**). RBD-specific IgG ELISA titer in the VibR group is relatively higher as compared to PoV2/3/4 group (**Fig. 1d**). Nucleocapsid-specific antibody response was detected in most of the individuals in VibR group and it was significantly higher compared with PoV2/3/4 vaccinated individuals (**Fig. 1e**). Together this data demonstrates the moderate elevation of antibody levels in individuals who tested positive for SARS-CoV-2 after vaccination compared to their previous samples.

Omicron-specific neutralizing antibodies are reduced after 6 months of vaccination and even after omicron infection, levels of neutralizing antibody levels against this variant were less^14^. We measured neutralizing antibodies before and after infection with the SARS-CoV-2 virus using a neutralization antibody competitive assay. We took a different approach instead of infection with real VOC, we used an assay which is based on the basic principle that the neutralization antibody circulating in the blood can bind to RBD region and block the binding of RBD and ACE2 which reduces viral entry. We quantified these functional neutralizing antibodies against different VOCs using LEGENDplex SARS-CoV-2 variants neutralizing antibody panel (5-plex) which is multiplex beads-based competitive assay. We examined the neutralizing antibody levels in PoV2 COVID-19 vaccinated individuals n=12 participants, PoV3 vaccinated individuals n= 6 participants and n=3 participant with PoV4 and VibR group (n= 21 participants). We divided the vaccinated individuals into two group either PoV2 and PoV3/4 and compared with the VibR group. Amongst PoV2 and PoV3/4 group an increasing amount of neutralizing antibodies (against all the known VOCs) was observed (**Fig. 2a**). When we compared the neutralizing antibodies data independently for PoV2, PoV3/4 and VibR groups, we could not observe a significant different among PoV3/4 and VibR groups. However, when data were compared with PoV2 and VibR groups we noticed significantly higher levels of neutralizing antibodies against previous VOCs (Fig. 2a). Further, we combined the all the vaccinated participants (PoV2/3/4) and compared the data with ViBbR group and found indeed overall, after infection neutralizing antibodies levels were upregulated against all the variants except for delta which did not reach to a significant level (**Fig. 2b)**. Overall, our results demonstrate that SARS-CoV-2 infection after vaccination boosts the neutralizing antibody response compared to vaccination alone.

**Fig. 2.**
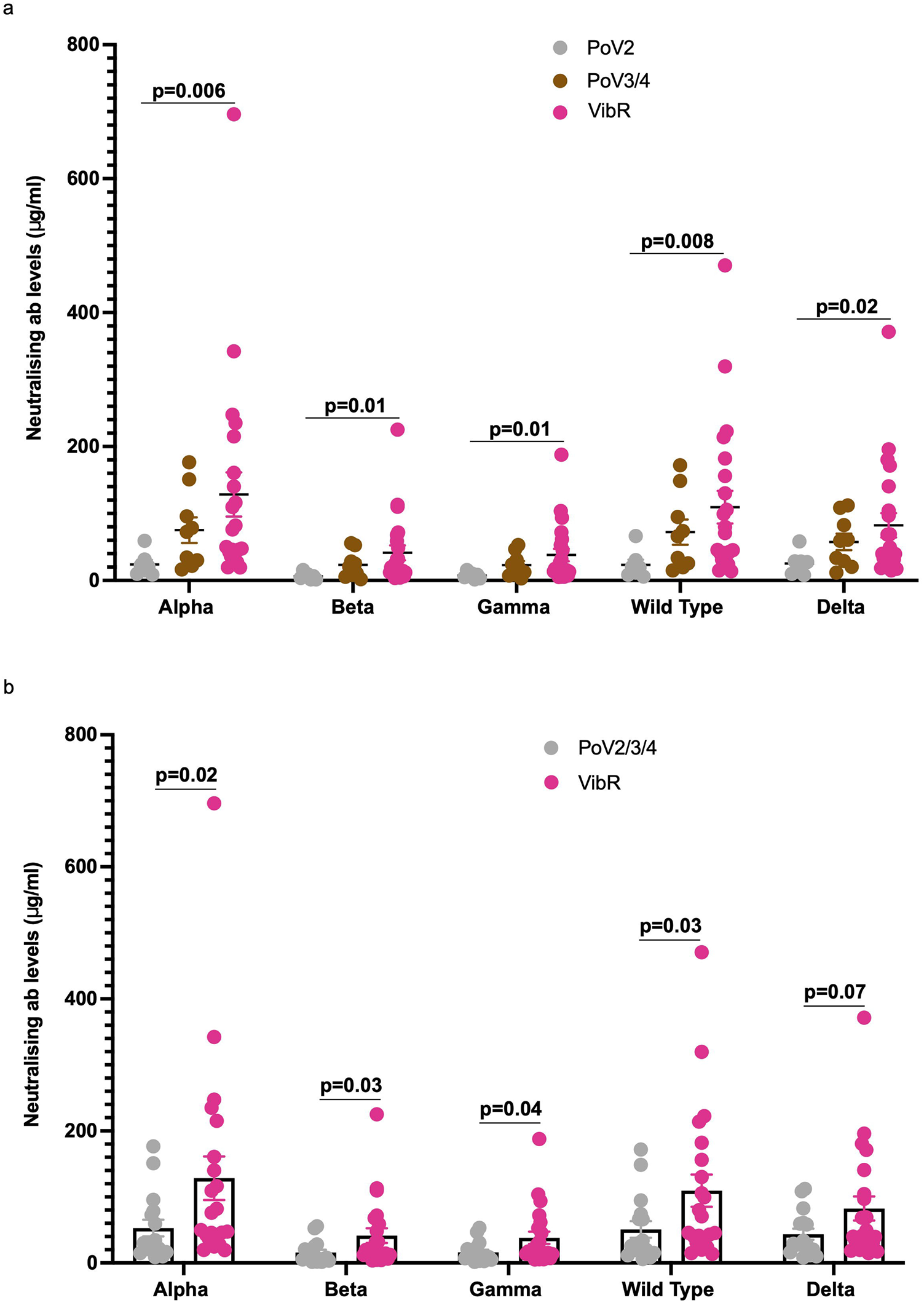
Neutralising antibody detection for different previous known variants of concern. (**a**) Neutralizing antibody levels after PoV2, PoV3/4 and VibR (after 14-30 days of recovery). (**b**). Neutralizing antibody levels against alpha, beta, gamma, wild type strain Wuhan and delta after PoV2/3/4 and VibR. Wilcoxon rank-sum test (unpaired) for two sample comparisons for PoV2/3/4 vs VibR and PoV2 vs VibR groups. Violin plots indicate mean and frequency distribution of the data including the interquartile range.

### Understanding the cellular immune response in VibR samples using dimensional reduction method

To understand the cellular immune response at single cell protein level, we employed 40 individual antibodies labelled with specific-fluorochromes to detect by spectral flow cytometry. The PBMCs from vaccinated (PoV2/3/4) and VibR samples were stained with a 40-colour antibody panel. Dead cells and doublets were removed in the gating strategy (**Suppl. Fig. 1**) and the data were combined (described in full in the material and methods) for high dimensional reduction method analysis. Using uniform manifold projection and approximation (UMAP) and Louvain unsupervised clustering analysis, we identified different immune clusters based on protein expression (**Fig. 3a**). Individual markers are shown for all the cell clusters (**Fig. 3b**). First, we characterized lymphocytes and monocytes which are separated into different major cell types including CD4^+^ T cells, CD8^+^ T cells, NKT cells, CD3^-^ HLA-DR^-^ CD56^+^ NK cells, ILCs, CD19^+^ B cells and CD4^+^ regulatory T cells (Tregs) (**Suppl. Fig. 1a-e**). To verify each cluster, we performed supervised clustering based on monocyte and lymphocyte gating (**Fig. 3b**) followed by cell annotations for individual key markers for specific cell types (**Fig. 3c, Suppl. Fig. 1a-e**). In individual antibody protein plots, we have observed the expression of individual markers such as CD45, which is ubiquitously present in cells such as lymphocytes and monocytes but not in myeloid cells. CD3 CD8, CD4, CD127, CD27, are highly expressed in the lymphocyte population. CXCR5 is highly expressed in CD19^+^ B cells, which is a marker for T follicular helper cells. PD-1 is a marker for exhaustion, expressed in many cell types like CD8^+^ T cells, CD4^+^ Tregs cells, and in monocytes (**Fig. 3c**). CD11c and CD14 is expressed in monocyte populations whereas HLA-DR is expressed by monocyte and B cell populations (**Fig. 3c, Suppl. Fig. 1**). Overall, we identified major immune cell subsets using unsupervised clustering and supervised cell annotations.

**Fig. 3.**
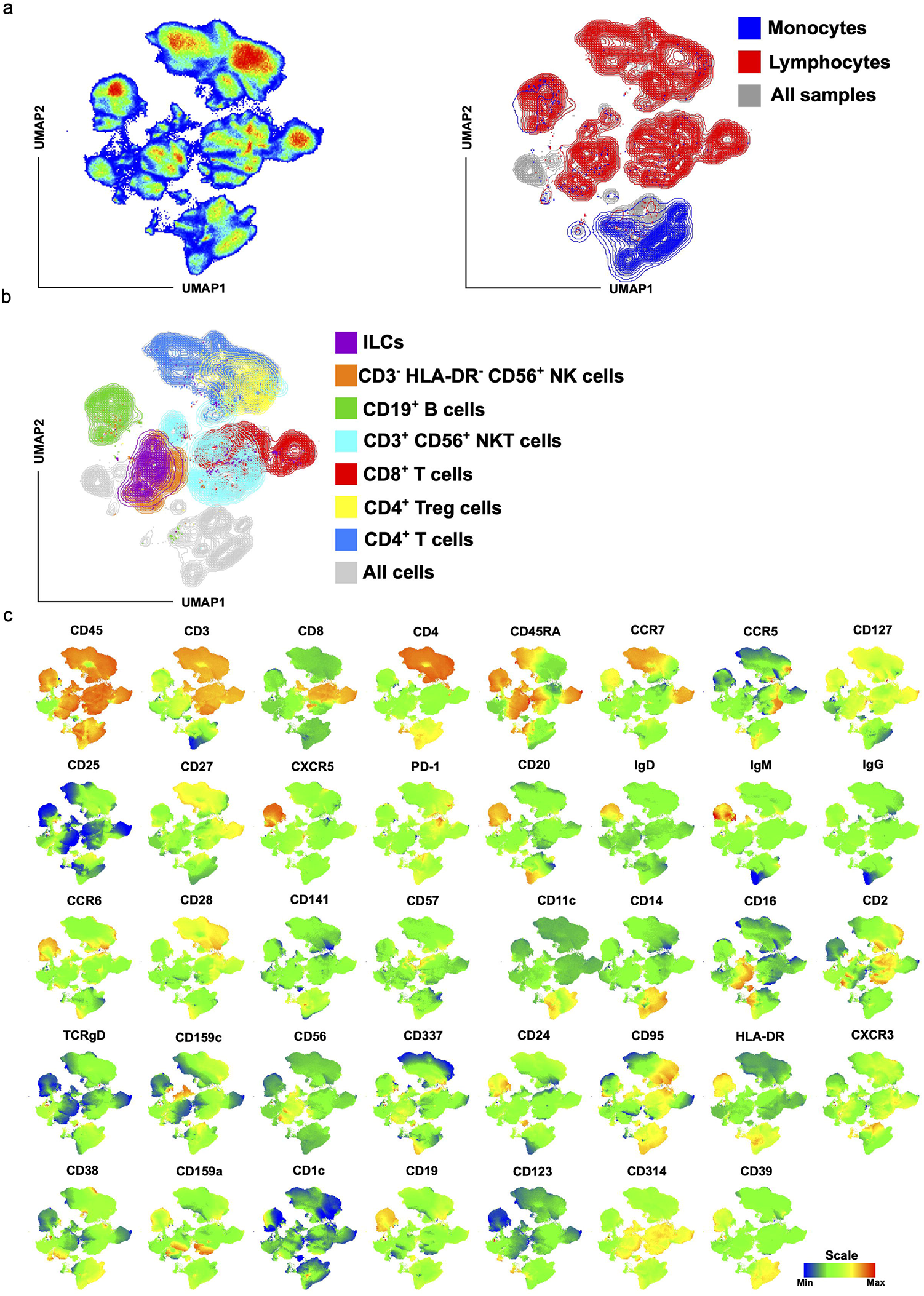
UMAP analysis for different cell subsets based on the expression of specific markers. (**a**) UMAP plot depicts monocytes and lymphocytes populations in participants samples. (**b**) UMAP plot representing lymphocyte subpopulations like ILCs, CD4^+^ T cells, CD8^+^ T cells, NK cells, NKT cells, and CD4^+^ T reg cells. (**c**) UMAP plots for individual antibody shows the expression of specific cell markers on specific cell populations like CD45, CD3, PD-1, CCR7, CD8, CD28, CD314, CD95.

### Dysregulated cellular immunity after infection in VibR individuals

CD3^-^CD19^-^CD20^-^ myeloid cells were decrease in percentage VibR group compared with PoV2/3/4 (**Fig. 4a-b**). For cell clusters like CD19^+^ CD20^+^ B cells, there was also increase in cell population in VibR group compared with PoV2/3/4 group. An opposite trend has been seen in CD45^+^/lymphocytes, NK cells in VibR participants. Tregs population was not affected after infection (VibR group) compared with vaccination individuals (**Fig. 4b**). Additionally, there was decline of the lymphocyte population in VibR group, in unpaired comparison or pairwise respectively (**Fig. 4c**). Next, monocytes were significantly upregulated based unpaired comparison or pairwise respectively in VibR group compared with vaccinated individuals (PoV2/3/4) (**Fig. 4d)**. The CD3^-^ CD19^-^ HLA-DR^-^ CD56^+,^ population was defined as NK cells. NK cells were significantly reduced in VibR group compared with vaccinated (PoV2/3/4) group (**Fig. 4e**). Furthermore, exploring the percentage of the CD3^-^ CD19^+^ CD20^+^ B cell population, higher levels of were found in VibR group compared to PoV2/3/4 participants (**Fig. 4f**). The machine learning workflow Tracking Responders EXpanding (T-REX) algorithm can capture phenotypic regions where significant change can occur between a pair of samples from one individual upon SARS-Cov 2 infection^20^. We, therefore, performed T-REX analysis on vaccinated (PoV2/3/4) and VibR groups to understand the expansion of specific immune cells after infection. Importantly, we find an expansion of CD4^+^ T cells, CD8^+^ T cells, B cells, monocytes, myeloid cells whilst decrease in NK cells population in most of the delta/omicron recovered individuals (VibR) (**Suppl. Fig. 2**).

**Fig. 4.**
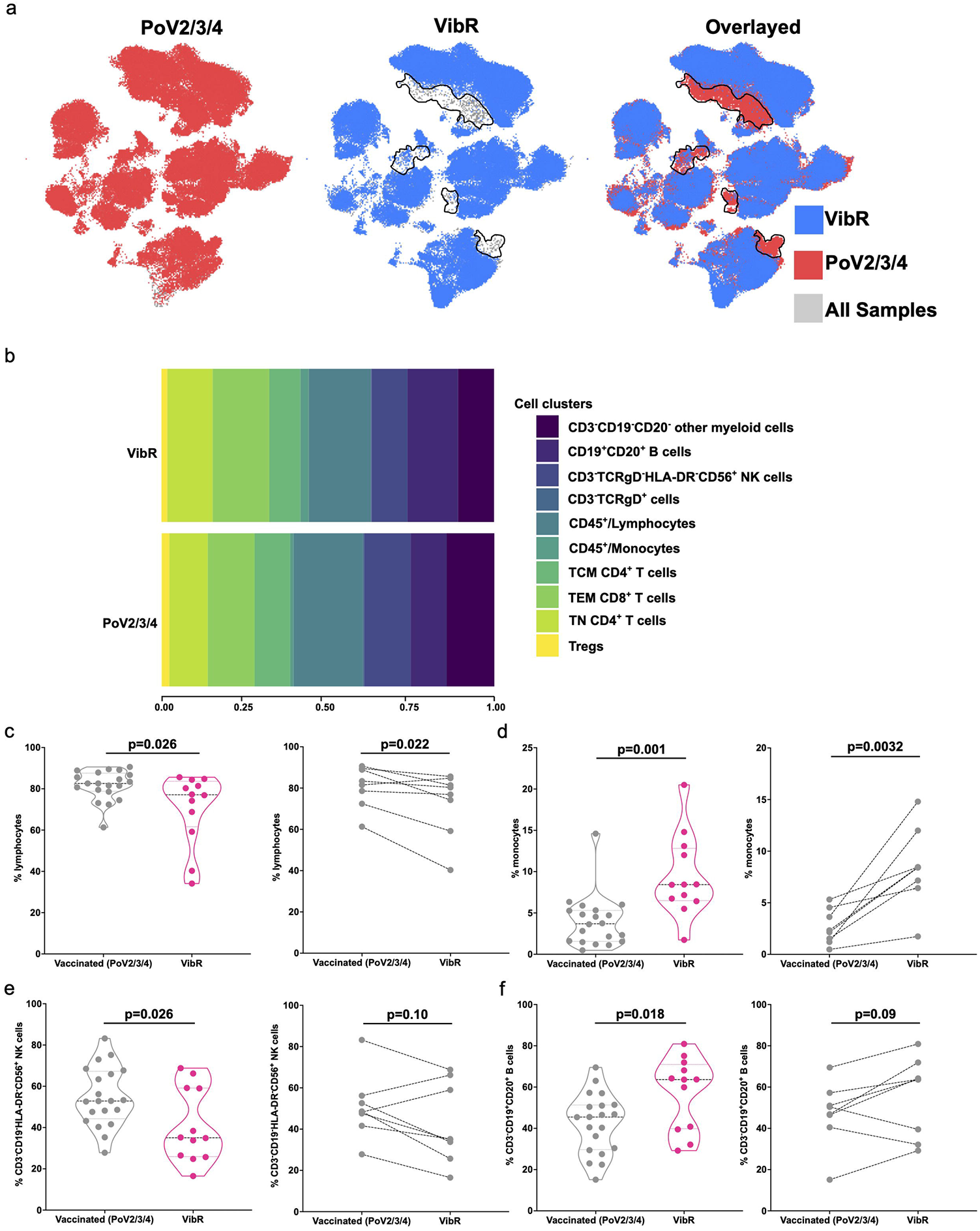
SARS-CoV2-delta/omicron-specific immune response on the expression of different markers before and after vaccination or infection. (**a**) Different clusters depict the cell population before and after infection. The right panel shows the difference in some cell types when overlayed. (**b**) Cell clusters like myeloid cells, B cells, NK cells, -TCR^+^ cells, lymphocytes, monocytes, CD4^+^, CD8^+^ T cells and γδ regulatory T cells before and after vaccination or infection. (**c**) Percentage of lymphocytes before (PoV2/3/4) and after infection (VibR) in vaccinated individuals. Lymphocytes were significantly reduced (p= 0.026; Wilcoxon rank-sum test) in infected VIbR group. Violin plots indicate mean and frequency distribution of the data including the interquartile range. The right panel (violin plot) represents all the samples (n=32) and left panel (paired dot plot) shows only paired samples from each individuals (n=8). (**d**) Percentage of monocytes before and after infection. The left panel represents that all infected individuals show a significant (p=0.001; Wilcoxon rank-sum test) increase in monocyte populations after infection, when monocytes were compared for the paired samples, a significant (p=0.0032; Student T-test) increase in monocytes were observed. The right panel (violin plot) represents all the samples (n=32) and left panel (paired dot plot) shows only paired samples from each individuals (n=8). (**e**) CD3^-^ CD19^-^ HLA-DR^-^ CD56^+^ NK cells were significantly reduced (p=0.02; Wilcoxon rank-sum test) in VibR group compared with PoV2/3/4 group (left; violin plot). However, a reduced population of NK cells was observed when pairwise comparisons were made among VibR group compared with PoV2/3/4 group (right; dot plot). Violin plots indicate mean and frequency distribution of the data including the interquartile range. (**f**) B cell populations (CD3^-^ CD19^-^ CD20^-^) was significantly increased (p=0.018; Wilcoxon rank-sum test) in VibR group compared with PoV2/3/4 group (left; violin plot). However, an increased population of B cells was observed when pairwise comparisons (p=0.09, Student T-test) were made among VibR group compared with PoV2/3/4 group (right; dot plot). Violin plots indicate mean and frequency distribution of the data including the interquartile range.

## Discussion

Previous studies revealed the significance of cellular and humoral immune response for sensitivity to COVID-19^3^. Our study aims the importance of adaptive immune response induced by vaccination with mRNA/DNA-based vaccines even after breakthrough infections and reveal that vaccine induced T cell response is sufficient to provide long-term protection. Here, we demonstrate the persistence of humoral immune response in COVID-19 vaccination (2-4^th^ doses) infected recovered participants compared to the vaccinated individuals using 40 colour flow cytometry methods. We found that vaccinated participants recovered from infection have higher spike specific IgG and RBD antibody levels to a vaccinated “only” group (2-4^th^ doses). We confirmed the infection in the recovered individuals by performing nucleocapsid specific IgG antibody assay, where post vaccination infected recovered individuals show higher nucleocapsid specific antibody levels. Previously, studies suggested that after 2^nd^ dose, omicron-neutralizing titers were reduced up to 22-fold compared with Wuhan-neutralizing titers^21^. Same study also described that one month after the 3^rd^ vaccine dose, omicron-neutralizing titers were increased 23-fold relative to their levels after two doses and were similar to levels of Wuhan-neutralizing titers after two doses^21^ and speculated that the requirement of a 3^rd^ vaccine dose of the mRNA vaccine BNT162b2 may protect against omicron-mediated COVID-19. In our study, we found that most of neutralizing antibodies against five different variants (WT (Wuhan), alpha, beta, gamma and delta) increased after 3^rd^ dose as well as further augmented after infection. Further, neutralizing antibody-response against different variants show that after the delta/omicron wave, vaccinated individuals have high number of neutralizing antibodies against WT (Wuhan), alpha, beta, gamma, and delta.

Cellular immune response was activated by vaccination (2-dose regime) and also by 1^st^ or 2^nd^ booster doses (3^rd^ or 4^th^ doses) and found to have antigen-specific T cell response (Singh Y et al., unpublished and published studies^22,23^). In case of triple vaccinated individuals, B and T cell immunity against previous VOCs was enhanced^24^. Our data are also in agreement with this finding. We showed that delta/omicron infection led to reduced proportions of T lymphocytes and monocytes as original Wuhan strain did to our immune response and this is supported by previously published findings^25,26^. Recent studies highlighted those infections reduced T cell reactivity and led to less neutralizing antibodies against B.1.1.529 (omicron) in naïve infected individuals, whereas in case of triple vaccinated individuals’ magnitude of T cell response was reduced ^24^. However, further experiments are required to validate these findings. NK cells appeared to have a key role in modulation of viral infection and these cells were reduced after infection in recovered individuals after 3^rd^ or 4^th^ vaccination. Taken together, these results show the markedly higher B cell population, but reduced monocytes and CD4^+^ T cells, which are required to provide effective humoral immune response to prevent from the severity of infection and to fight against COVID-19. In sum, our study provides a baseline for future immunophenotyping to measure infected recovered individuals. This approach could be applied to understand the longitudinal immune response in COVID-19.

### Limitations

Our study has a number of limitations; the primary limitation is the collection of samples to perform in depth analysis of immune response. We have only 14 donors in this whole study, and for some of the individuals we have had blood samples only from the post vaccination infection stage as they did not participate earlier in COVID-19 VAC program. Additionally, after the infection, white blood cells are generally low, so, it was incredibly difficult to get the enough number of cells for performing antigen specific T cell response.

## Figure legends

**Suppl. Fig. 1 Gating strategy for 40-colour immunophenotyping**

A 40-colour antibody and live/dead staining was performed as described in material and methods. (**a**) First from individual FCS file, debris removal gate was applied to keep the live cells. These live cells were identified using Live/dead staining dye. Afterwards, singlets were identified using FSC-A vs FSC-H and singlets were gated for CD45+ lymphocyte population. CD45+ lymphocytes were again represented in FSC-A vs SCC-A to identify the lymphocyte and monocytes based on size and granularity. (**b**) Lymphocytes were separated into three major populations based on CD3 and γδ-TCR antibodies for CD3^+^ T cells, γδ-TCR^+^ cells, and CD3^-^ γδ^-^ cells. CD3^+^ γδ-TCR^-^ cells were used for identification of NKT cell population based on CD3^+^ CD56^+^ NKT cells as in FACS panel (CD56 vs CD8). CD3^+^CD56^-^ T cell CD5 population was divided into CD4 and CD8 population (CD4 vs CD8). Individual CD4 or CD8 cell population was identified for naïve, effector memory, central memory, TEMRA and Tregs cell populations using CCR7, CD45RA, CD27, CD28, CD127, CD25, CD39 markers as described in the FACS plots. (**c**) Identification of monocytes (**d**) B cells (**e**) and NK cell subsets as described in FACS plots for a specific cell population.

**Suppl. Fig. 2Antigen-specific expansion of T cells using TREX analysis**.

Total of 8 individuals were analysed before and after recovery with delta/omicron infection. UMAP plots show expression of individual markers (left) for 19 different immune cell markers including T, B, NK, ILCs, γδ-T cells, monocytes, and dendritic γδ cells. Expansion of antigen-experienced T and B cells after infection (right panel; orange).

## Materials and Methods

### Human subjects and experimentation

All the enrolled vaccinated infected but recovered (VibR) participated in our earlier COVID-19 VAC study. Total 32 participants (25 out of 110 (COVID-19-VAC participants) VibR participants infected with either with delta or omicron VOC SARS-CoV-2 during December 2021 – April 2022) were enrolled in this study. Additional, 7 VibR participant also participated in this study for whom we did not have pre-infection PBMCs/plasma samples. All the vaccinated individuals were either vaccinated 2 times with recombinant Adenoviral vector (rAdVV) or mRNA (ChAdOx1 nCoV-19; AstraZeneca (AZ), Ad26.CoV2.S; Johnson & Johnson (JJ), BNT162b1; Pfizer/BioNTech (PB), mRNA-1273; Moderna (MD)) and have had either 3^rd^ or 4^th^ doses (1^st^ or 2^nd^ booster with mRNA (BNT162b1; Pfizer/BioNTech (PB), mRNA-1273; Moderna (MD)) of COVID-19 vaccines. All the volunteers were recruited for the study under informed consent and a baseline health questionnaire was also completed. We did not provide the vaccine in our Institute and volunteers were free to choose to take the vaccines as per guidelines of the state or federal governments. No ethnic data were collected, however, most of the participants were from university town Tübingen area with multinational background. The study was approved by the University Hospital Tübingen, Eberhard-Karls University of Tübingen, Institutional Review Board (Ethik-Kommission an der Medizinischen Fakultät der Eberhard-Karls-Universität and am Universitätsklinikum Tübingen, Germany, Ethics no: 355/2021BO2 (Clinical Trial.gov-Nr.: NCT04873128) and 286/2020BO1 Clinical Trial.gov-Nr.: NCT04364828). The study was conducted within full compliance of Good Clinical Practice as per the Code of German Regulation. Based on epidemiology and genetic testing studies (based on Robert Koch Institute, Germany), therefore, most of vaccinated individuals have infection either with delta or omicron variants of SARS-CoV-2 during the sample.

### PBMC and Plasma isolation

Whole blood was collected in 9-ml NH4^+^-Heparin tubes, the upper layer was transferred into Eppendorf tubes and blood samples were processed with in 2 hours of blood collection for PBMC and plasma isolation protocol. Blood tubes were allowed to stand in upright position until used or at least for 30 minutes and 1ml of upper layer was collected and transferred to 2.0 mL Eppendorf tubes. Eppendorf tubes were centrifuged at 2000 x g for 10 mins at RT. Supernatant (Plasma) was transferred into cryovials and stored at -80°C for further experiments. PBMCs were isolated from blood collected in the heparin tubes by density gradient medium. First, in a sterile 50 ml falcon, 15 ml Pancoll was added and allow to rest the tube. In another 50 ml falcon tube, blood was gently mixed by inverting the tube with 1:2 ratio of blood and 1xPBS. Diluted blood was gently mixed with 25 ml pipette for a homogenous mixing of blood and PBS. After mixing, diluted blood was layered on a density gradient (Pancoll), and centrifuge at 400 x g for 22 mins (without breaks; 1 acceleration and 0 deceleration) at room temperate (RT). All steps were performed at RT. Buffy coat was collected with a pipette and added 20 ml 1x PBS. PBMC were washed 2 times in 1xPBS and subsequently frozen in freezing media [70 % fetal bovine serum (FBS) with 30% dimethyl sulfoxide (DMSO)] at -80 °C in Corning™ CoolCell™ LX Cell Freezing Vial Containers for overnight then vials were transferred to normal boxes for storage in – 80 °C freezer.

### Anti-spike protein S1, anti-spike protein RBD and anti-Nucleocapsid IgG antibodies multiplex flow cytometry ELISA

A 3-plex SARS-CoV-2 Serological IgG Panel (BioLegend LEGENDplex, Cat. No. #741132) was used for measuring IgG titre of the 3 key COVID-19 coronavirus derived proteins: spike1 (S1), receptor binding domain (RBD) and nucleocapsid (N) and measured the antibody levels as per manufacture’s recommendation. Standards were prepared by serial dilutions (1:4) and top standard contained the highest concentration (10^5^ ng/mL). Plasma samples were diluted 800-fold with assay buffer. V-bottom plates were used to perform the assay. Standards and samples were run in duplicates. 25μL of each assay buffer and standards or samples were added to each well. Then 25μL of capture beads, which can bind to IgG of S1, RBD and N, were added to each well. The plate was shaken at 800 rpm for 2 hours at RT. Beads were then spun down (1050 rpm for 5 minutes) and supernatant was removed. After 1 time washing by using 1X Wash Buffer, 25μL of biotinylated detection antibody was added to each well, thus capture bead-IgG-detection antibody sandwiches were formed. The plate was then shaken at 800 rpm for 1 hour at RT. Streptavidin-phycoerythrin (SA-PE) was subsequently added without washing, which can bind to the biotinylated detection antibodies and provide fluorescent signal. Plate was then shaken at 800 rpm for 30 minutes at RT. Supernatant was removed by centrifugation (1050 rpm for 5 minutes), followed by one time washing. After resuspending the beads, PE signal fluorescence intensity of all the samples was then quantified by using flow cytometry.

### Neutralizing 5-plex antibody assay

A 5-plex SARS-CoV-2 Variants Neutralizing Antibody Panel (BioLegend LEGENDplex, Cat. No. #741174) was used to measure the neutralization antibody concentration. Capture beads contains five SARS-CoV-2 spike protein variants (variant Alpha, variant Beta, variant Gamma, Wild type S1 protein and variant Delta) conjugated onto fluorochrome-coded beads. Biotinylated ACE2 protein serving as the detection which was added to all wells after adding the beads. Standard (anti-human S1 RBD recombinant antibody) or samples and detection competed for binding to the capture beads, producing a competitive assay. Streptavidin-phycoerythrin (SA-PE) was subsequently added, which bind to the biotinylated detection, providing fluorescent signal intensities in proportion to the amount of bound analytes and less neutralizing antibodies for the corresponding VOC. Standard solutions were prepared by serial 1:3 dilutions of top standard (10^5^ ng/mL, Anti-human S1 RBD recombinant antibody). Plasma samples were diluted 100-fold with Assay Buffer (2μL sample + 198μL Assay Buffer). The assay was performed in a 96-well V-bottom plate and standards were run in duplicate. After adding samples or standards and Assay Buffer, 25μL of beads (SARS-CoV-2 Spike protein Variants: Variant Alpha, Beta, Gamma and Delta, and Wild type S1 protein) were added to each well. Biotinylated ACE2 protein serving as the Detection, which will compete with Standards or samples to bind to the beads, were then added to all wells. Plate was shaken at 800rpm for 2 hours at room temperature (RT). 25μL of Streptavidin-phycoerythrin (SA-PE) were subsequently added, which will bind to the bound biotinylated ACE2 protein, providing fluorescent signal intensities in proportion to the amount of bound analytes. The plate was then shaken at 800 rpm for 30 minutes at RT. Beads were spun down (1050 rpm for 5 minutes) and supernatant was removed. After 2 times washing, beads were resuspended by 150μl Wash Buffer. Measurements were performed by using flow cytometer.

### Flow cytometry staining

First, antibody cocktail has been made by adding the antibodies mentioned in the Table 1 and kept at 4°C in the dark. Then, prepared the 50 KU/DNase I/FACS buffer by adding 1.91 μl DNase I in per ml FACS buffer and 200 KU DNase I /FACS buffer by adding 7.64 μl DNase I per ml FACS buffer was prepared.

In vitro stimulation of freshly isolated and frozen PBMC stained with specific cell surface markers was performed. In this assay, we used 1 - 3 million cells for staining purpose. Frozen PBMC have been thawed at 37°C for 1-2 mins (2 samples at one time). PBMCs were then transferred in 10 ml 50KU DNase I/FACS Buffer followed by the centrifugation for 10 mins at 350xg at RT. After every centrifugation step, the supernatant was discarded. Cell pellet was resuspended in 5 mL of 200KU DNase I/FACS buffer counted the cells before the staining. Samples were incubated for 20 mins at 4°C in the dark, followed by centrifugation for 10 mins at 350xg. Cell pellet was resuspended in 100 -150 μl 50KU DNase I/ FACS buffer. 3 million cells or all sample was added into the 96 well plate (U-bottom) and centrifuged for 10 mins at 350xg at 4°C. Afterwards the sample was incubated with 150 μL human IgG at least for 20 mins at 4°C in the dark. After incubation, plate was centrifuged at 350xg for 5 mins at 4°C. Live/dead staining was performed by using viability dye (1:4000) followed by the incubation for 30 mins at RT. Further, 2 washing steps have been performed by using 50KU DNase I/ FACS buffer. Afterwards 50 μL Brilliant stain buffer and 5 μL Monocyte blocking reagent were added to each sample before 100 μL antibody cocktail were added to each sample. Samples were mixed well and incubated for 15 mins at 37 °C, followed by 30 mins incubation at RT in the dark. Afterwards, 50 μL 50KU DNase I/ FACS buffer was added and samples were centrifuged for 5 mins at 350xg and 4°C. Washing step was repeated with 200 μL 50KU DNase I/FACS buffer and cell pallet was resuspended again in 200 μL 50 KU FACS buffer. Sample was transferred into FACS tube and kept the sample on ice until run on a flow cytometer (Cytek). Samples were run within few hours of staining on the same day as no fixation steps has been performed. At least 0.5 million cells were acquired at the CYTEK Aurora.

### Flow cytometry data analysis

#### Normal percentage cell analysis

Vaccinated and vaccinated recovered after delta/omicron infection samples were processed. First FSC-H vs SSC-H cells were gated for debris removal and these gated cells live/dead staining vs SSC-H to remove the dead cells. Live gated cells were plotted for FSC-A vs FSC-H to get the singlet cells. Further singlet cells were gated for CD45^+^ lymphocytes and then finally investigated for monocytes and lymphocytes. All the different T, NK, B, monocytes, ILC, gdT cells were analysed as described earlier^27^.

#### Dimension reduction analysis using uniform manifold approximation and project (UMAP)

All the FACS files were concatenated and gated as normal percentage cell analysis and CD45^+^ lymphocytes were taken (50,000 cells from each sample). Concatenated files were subjected to 39 colour UMAP analysis and manually annotated based on individual marker expression.

#### T-REX Data analysis

After normalization, FACS data were scaled with an arcsinh transformation, with an appropriate cofactor set for each channel following standard procedures for fluorescence data as described earlier^20^. The FACS data were then manually gated for removal of atypical events. After quality control gating, a UMAP analysis was performed on the cleaned-up samples using the surface markers in each panel. The resulting common, 2-D embedding of the data was used for visualization and selection of either CD3^+^ T cells (from T cell panel data) or B cells (from B cell panel data) for further downstream analysis. A common t-SNE analysis was done on all 8 donor samples using the same markers used to create the UMAP on either the CD3^+^ T cells or B cells extracted from their respective UMAP. After the t-SNE, equally sampling was done on each donor pair.

### Data Analysis and statistics

Group comparisons were performed using GraphPad Prism version 9.0. Populations were compared using Mann-Whitney unpaired test rank sum tests. P values less than 0.05 were considered statistically significant.

## Acknowledgements

We are grateful to all the volunteers for donating blood samples for the study. We thank Dr Denis Witt, Dr Lisa Ruisinger, Dr Jeannette Huebner-Schmid and Miss Marion Loitz for technical assistance with blood sample collection and PBMCs isolation. Prof Dr Sara Y Brucker for infrastructure support for the experiments. We also acknowledge UKT, FACS core facility for providing excellent infrastructure for flow cytometry measurements and technical assistance. We gratefully acknowledge Prof Dr Huu Phuc Nguyen, Ruhr-Universität Bochum, Germany for critical reading of the manuscript.

## Funding

This project was supported by the Deutsche Forschungsgemeinschaft (DFG) to OR. YS was supported by Ferring Pharmaceutical and Fortüne funds (2642-0-0).

## Author’s contribution

RB: Performed PBMCs isolation, flow cytometry staining, antibody assays and data analysis

ZY: Performed the PBMCs isolation and antibody assays and flow cytometry staining

VH: Machine learning tools implementation (TREX) and data analysis

SP, KB: Flow cytometry assay and data analysis

MSS, NC, SO and OR: Experimental planning, data discussion, funding acquisition and manuscript editing

YS: Conception of the project, overall project management, data analysis and writing the manuscript

## Conflicts of Interest

The authors declare no conflict of interest. The funders had no role in the design of the study; in the collection, analyses, or interpretation of data; in the writing of the manuscript; or in the decision to publish the results.

## Data availability

FCS cell data are available from the corresponding author upon request.

## Notes

### Competing Interest Statement

The authors have declared no competing interest.

