## Supplementary material for "Immune dynamics at single cell protein level after delta/omicron infection in COVID-19 vaccinated convalescent individuals": Suppl. Info

Running title: COVID-19 infection in vaccinated individuals

### Suppl. Figures:

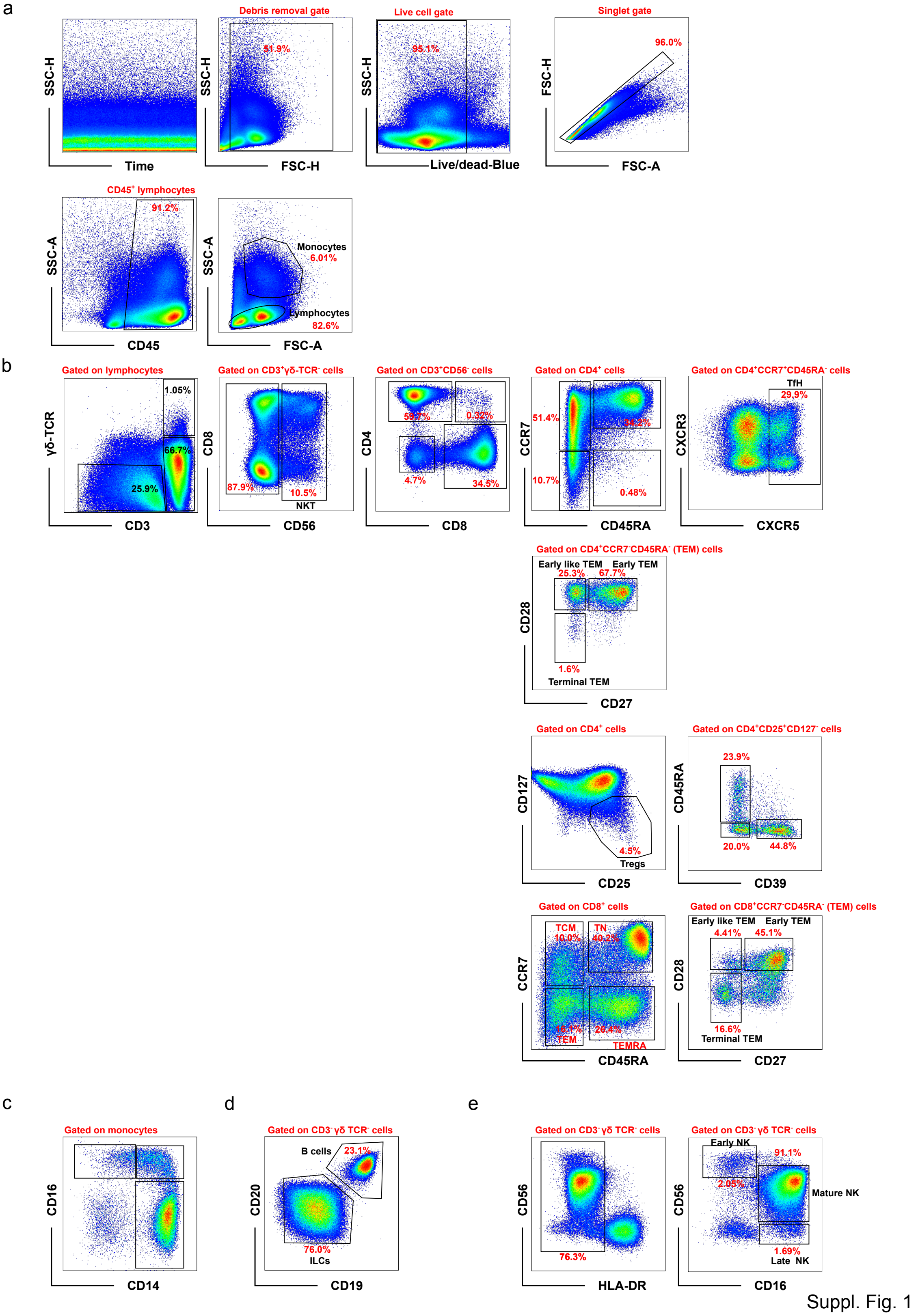


Suppl. Fig. 1

**Suppl. Fig. 1 Gating strategy for 40-colour immunophenotyping**

A 40-colour antibody and live/dead staining was performed as described in material and methods. (**a**) First from individual FCS file, debris removal gate was applied to keep the live cells. These live cells were identified using Live/dead staining dye. Afterwards, singlets were identified using FSC-A vs FSC-H and singlets were gated for CD45+ lymphocyte population. CD45+ lymphocytes were again represented in FSC-A vs SCC-A to identify the lymphocyte and monocytes based on size and granularity. (**b**) Lymphocytes were separated into three major populations based on CD3 and γδ-TCR antibodies for $CD3^+$ T cells, γδ-$TCR^+$ cells, and $CD3^-$ $\gamma\delta^-$ cells. $CD3^+$ γδ-$TCR^-$ cells were used for identification of NKT cell population based on $CD3^+CD56^+$ NKT cells as in FACS panel (CD56 vs CD8). $CD3^+CD56^-$ T cell population was divided into CD4 and CD8 population (CD4 vs CD8). Individual CD4 or CD8 cell population was identified for naïve, effector memory, central memory, TEMRA and Tregs cell populations using CCR7, CD45RA, CD27, CD28, CD127, CD25, CD39 markers as described in the FACS plots. (**c**) Identification of monocytes (**d**) B cells (**e**) and NK cell subsets as described in FACS plots for a specific cell population.

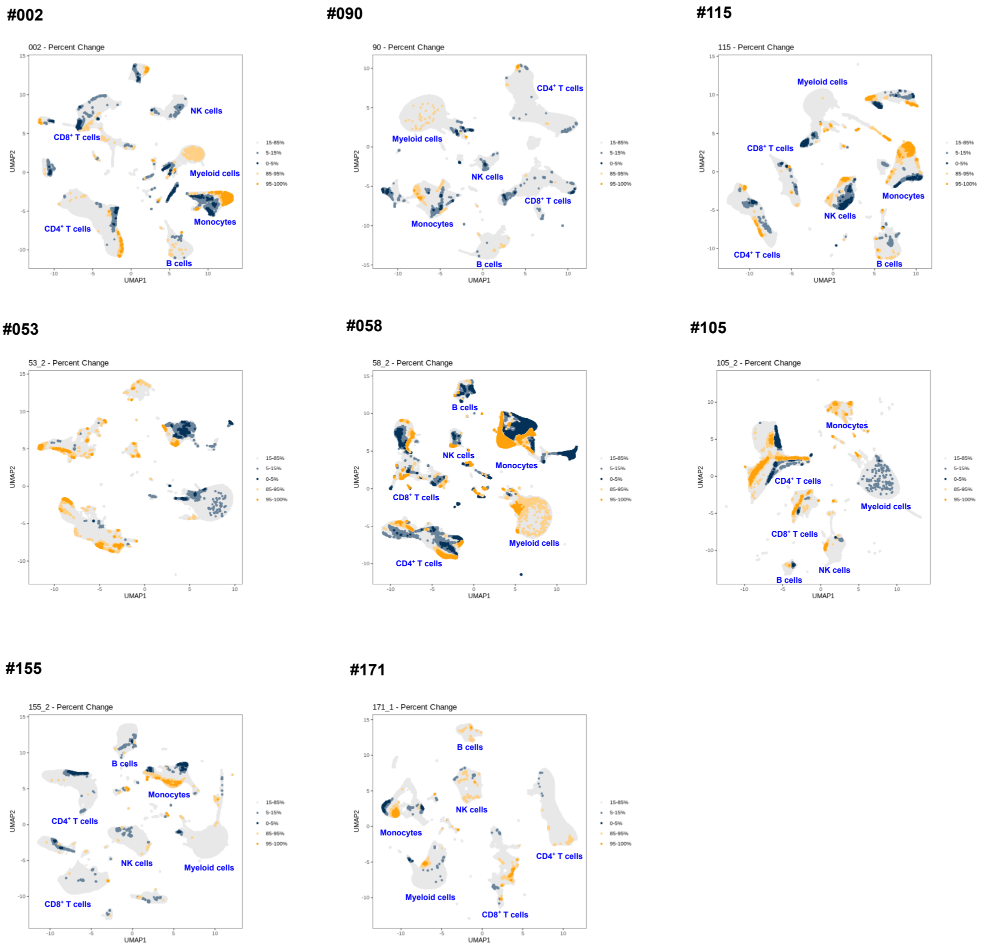


Suppl. Fig. 2

**Fig. 2 Antigen-specific expansion of T cells using TREX analysis.**

Total of 8 individuals were analysed before and after recovery with delta/omicron infection. UMAP plots show expression of individual markers (left) for 19 different immune cell markers including T, B, NK, ILCs, γδ-T cells, monocytes, and dendritic cells. Expansion of antigen-experienced T and B cells after infection (right panel; orange).
